# ERK hyperactivation in epidermal keratinocytes impairs intercellular adhesion and drives Grover disease pathology

**DOI:** 10.1101/2024.04.30.591953

**Authors:** Cory L. Simpson, Afua Tiwaa, Shivam A. Zaver, Christopher J. Johnson, Emily Y. Chu, Paul W. Harms, Johann E. Gudjonsson

## Abstract

Grover disease is an acquired dermatologic disorder characterized by pruritic vesicular and eroded skin lesions. While its pathologic features are well-defined, including impaired cohesion of epidermal keratinocytes, the etiology of Grover disease remains unclear and it lacks any FDA-approved therapy. Interestingly, drug-induced Grover disease occurs in patients treated with B-RAF inhibitors that can paradoxically activate C-RAF and the downstream kinase MEK. We recently identified hyperactivation of MEK and ERK as key drivers of Darier disease, which is histologically identical to Grover disease, supporting our hypothesis that they share a pathogenic mechanism. To model drug-induced Grover disease, we treated human keratinocytes with clinically utilized B-RAF inhibitors dabrafenib or vemurafenib and leveraged a fluorescent biosensor to confirm they activated ERK, which disrupted intercellular junctions and compromised keratinocyte sheet integrity. Consistent with clinical data showing concomitant MEK blockade prevents Grover disease in patients receiving B-RAF inhibitors, we found that MEK inhibition suppressed excess ERK activity to rescue cohesion of B-RAF-inhibited keratinocytes. Validating these results, we demonstrated ERK hyperactivation in skin biopsies of vemurafenib-induced Grover disease, but also in spontaneous Grover disease. In sum, our data define a pathogenic role for ERK hyperactivation in Grover disease and support MEK inhibition as a therapeutic strategy.

**GRAPHICAL ABSTRACT:** 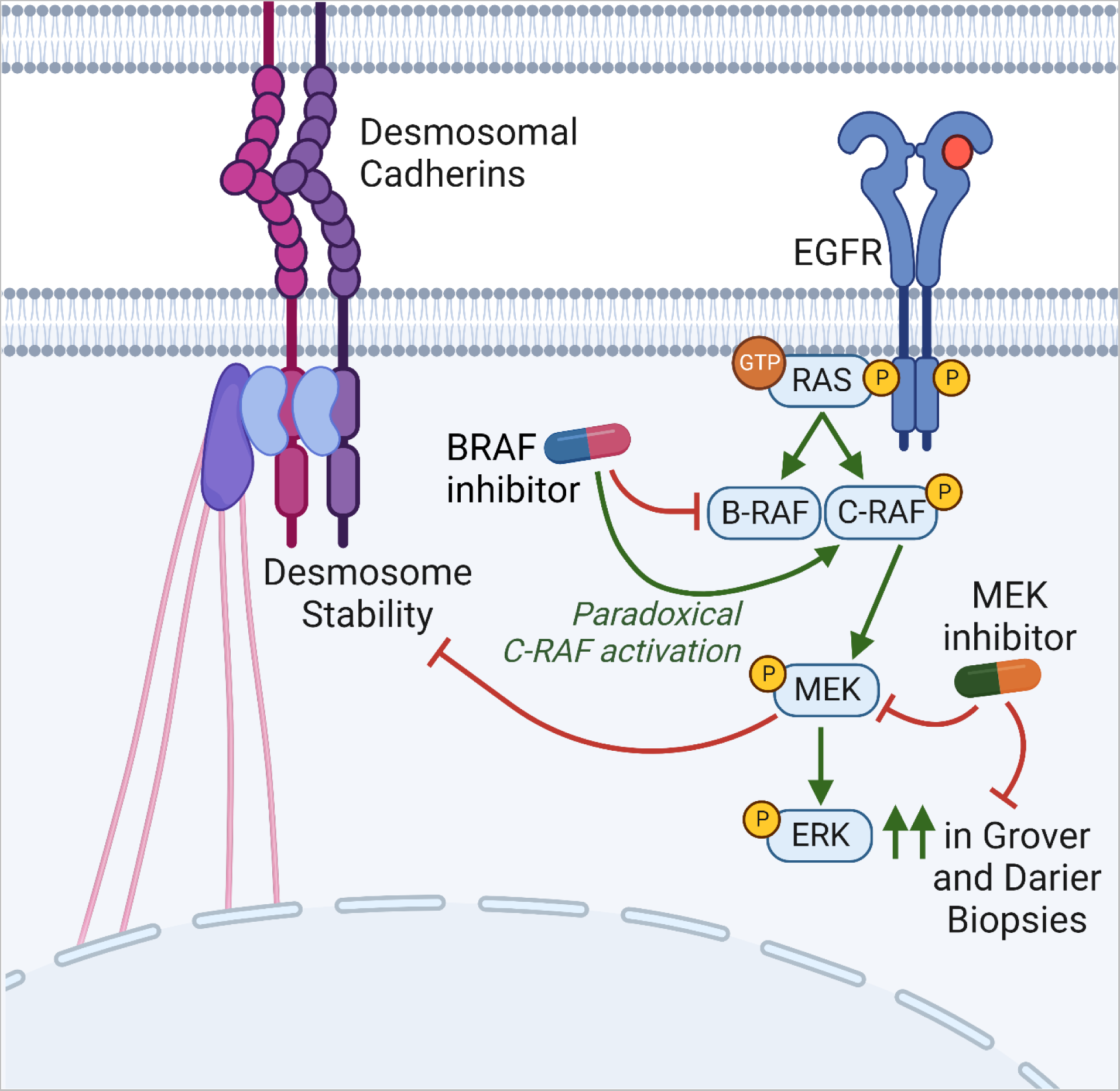

## INTRODUCTION

Grover disease (GD) is a spontaneous dermatologic disorder of unknown etiology that can cause intractable pruritus, extensive vesicular skin lesions in severe cases, and cutaneous superinfection (1-3). Since first described in 1970 (4), the pathological features of GD are well-defined as the combination of abnormal differentiation of keratinocytes (called dyskeratosis) and severing of their intercellular connections (termed acantholysis), which leads to blister formation within the epidermis (5, 6). However, the molecular drivers of these defects in epidermal integrity are poorly understood and there are no FDA-approved therapies for GD (7, 8). Interestingly, the pathologic features of GD can be indistinguishable from a genetic blistering disorder called Darier disease (9, 10), suggesting they may share a pathogenic mechanism. In fact, recent sequencing of GD lesions found acquired mutations in *ATP2A2* (11), the gene linked to DD (12, 13).

Both Grover- and Darier-like eruptions have been reported as common cutaneous toxicities of systemic therapy with B-RAF inhibitors including dabrafenib and vemurafenib (14, 15), which are utilized in the treatment of *BRAF* mutant cancers such as melanoma (16-19). Paradoxically, sustained B-RAF inhibition has been shown to induce *activation* of MEK (20, 21), which operates downstream of RAF in the mitogen-activated protein (MAP) kinase pathway, but how this signaling aberration induces the pathologic features of GD in human epidermal keratinocytes remains unknown. In recent work, we linked Darier disease to MEK and ERK overactivation (22), which led us to hypothesize that MEK hyperactivity might also fuel GD pathogenesis. Consistent with this, prior retrospective analysis of patients co-treated with MEK inhibitors, instead of B-RAF inhibitor monotherapy, failed to develop GD as a side effect (23).

Early studies of GD using histology and electron microscopy identified defects in desmosomes (6, 24), cell-cell junctions that are essential to maintain epidermal tissue integrity (25). Given numerous studies connecting the MAP kinase pathway to the stability of desmosomes (26-33), we proposed that overactivation of MEK and downstream ERK in GD could directly compromise intercellular adhesion. We modeled drug-induced GD in human keratinocytes and in 3D organotypic epidermis (34) using B-RAF inhibitors, which weakened cell-cell adhesion in an ERK-dependent manner. To validate a role for the MAP kinase pathway in GD pathogenesis, we assessed ERK activation in skin biopsies of B-RAF inhibitor-induced GD, which led us to propose MEK inhibition could also serve as a targeted therapy for *spontaneous* GD (**Graphical Abstract**).

## RESULTS

### Sustained B-RAF blockade paradoxically activates ERK in human epidermal keratinocytes

In patients treated chronically with systemic B-RAF inhibitors for malignancies driven by activating *BRAF* mutations, drug-induced GD is a common cutaneous toxicity (14, 15). This has been theorized to occur via off-target effects on bystander cells (like keratinocytes) that have wild-type *BRAF*. In these cells, B-RAF inhibitors cause compensatory upregulation of C-RAF, which leads to paradoxical *activation* of the MAP kinase pathway via downstream kinases MEK and ERK; this was originally shown in human anaplastic carcinoma cells (20). We treated normal human epidermal keratinocytes (NHEKs), the cells that manifest GD pathology, with either of the two B-RAF inhibitors most often reported to induce GD, dabrafenib (Dab) and vemurafenib (Vem). Treatment of keratinocytes with either B-RAF inhibitor was sufficient to increase the active phosphorylated form of ERK (pERK) compared to drug vehicle (dimethyl sulfoxide, DMSO) as quantified by Western blotting (WB) of NHEK lysates; ERK activation was dampened by using trametinib to inhibit MEK, the upstream kinase in the MAP kinase pathway (**Figure 1A-B**).

**Figure 1.**
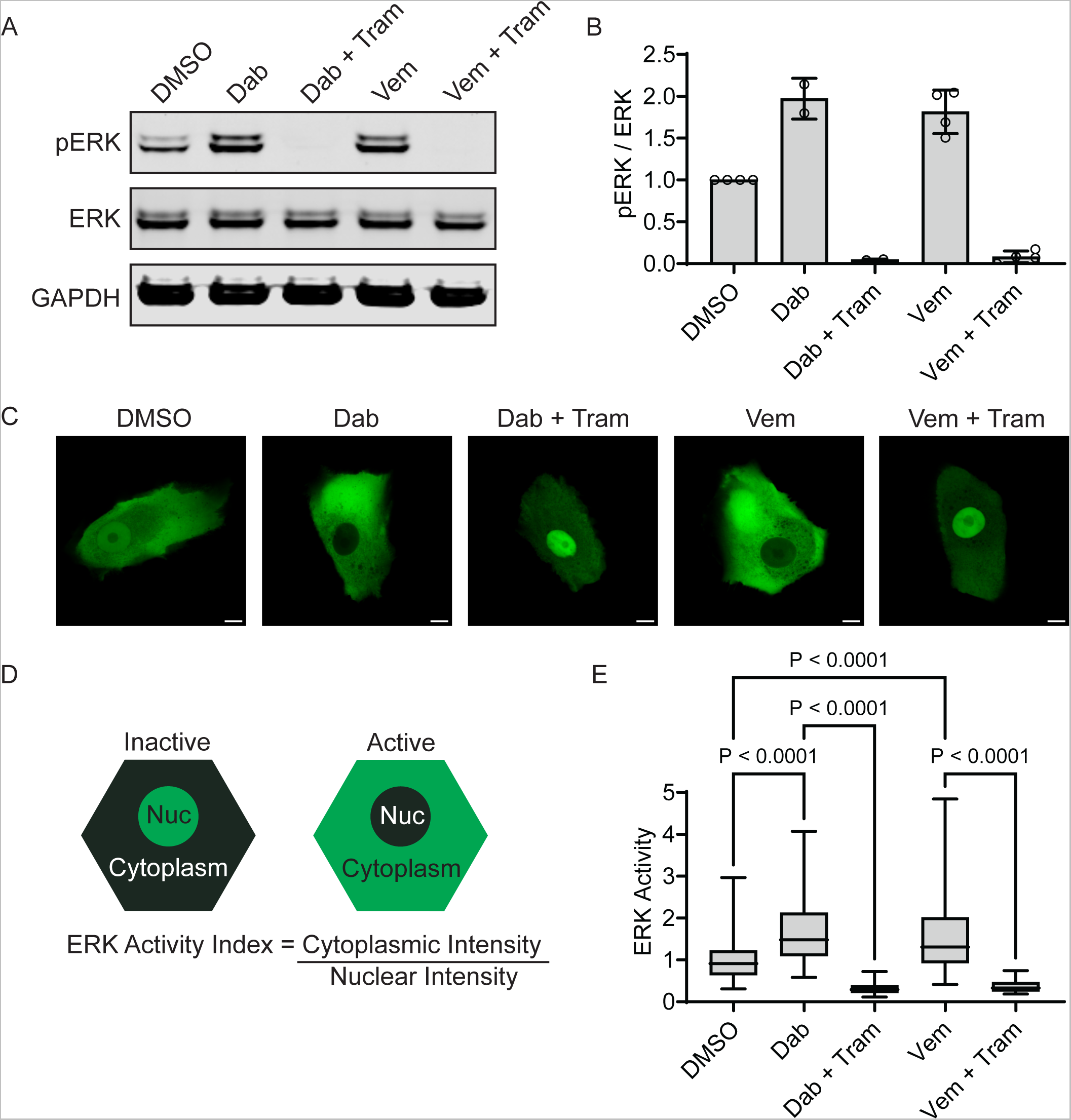
Sustained B-RAF blockade paradoxically activates ERK in human epidermal keratinocytes. **(A)** Immunoblot of total and phosphorylated ERK (pERK) in lysates from NHEKs treated with dabrafenib (Dab, 1 μM) or vemurafenib (Vem, 10 μM) +/-trametinib (Tram, 1 μM) for 24 h; GAPDH is a loading control. **(B)** Bar graphs display the mean ± SD of the intensity of pERK (normalized to total ERK) with individual data points plotted for N=2 (Dab) or N=4 (Vem) independent experiments. **(C)** Representative confocal fluorescence microscopy images of NHEKs transduced with the ERK biosensor (ERK-KTR) linked to the green mClover fluorophore; cells were treated with the indicated compounds for 24 h in medium containing 1.2 mM calcium; scale bar = 10 µm. **(D)** Diagram of the ERK biosensor, which is primarily localized in the nucleus when ERK is inactive vs. in the cytoplasm when ERK is active; an ERK activity index is calculated as the cytoplasmic to nuclear fluorescence intensity ratio. **(E)** ERK activity data for each treatment group are depicted as a box plot of the 25^th^-75^th^ percentile with a line at the median from N≥66 cells per condition from 2 biological replicates; mean ERK activity of DMSO was normalized to 1; P-values are from one-way ANOVA using the Bonferroni adjustment for multiple comparisons.

We substantiated these findings in live NHEKs using a fluorescent biosensor engineered to shuttle out of or into the nucleus upon ERK activation or inactivation, respectively (35). In live NHEKs transduced with the ERK kinase translocation reporter linked to a green fluorescent protein (ERK-KTR-Clover), we found that treatment with dabrafenib or vemurafenib induced ERK activation in a manner that could be reversed by inhibiting MEK (**Figure 1C-E**). These findings confirm that NHEKs exhibit paradoxical activation of ERK upon sustained treatment with selective B-RAF inhibitors previously linked to GD in clinical studies (14, 15).

### B-RAF inhibition disrupts desmosomal protein localization in epidermal keratinocytes

ERK is well known to regulate cell-cell adhesion and differentiation of keratinocytes (28, 30, 36, 37), but this MAP kinase has not been directly linked to GD pathogenesis. To test if B-RAF inhibitor-induced ERK activation impaired intercellular adhesion, we assessed the level and localization of cell-cell junction proteins in NHEKs. Compared to control keratinocytes treated with DMSO, the B-RAF inhibitors dabrafenib or vemurafenib did not appreciably alter protein levels of classical cadherins, desmosomal cadherins (DSG1, DSG2, DSG3) or their catenin binding partner plakoglobin (PG) (**Figure 2A**). However, using immunofluorescence (ImF) microscopy, we found both dabrafenib and vemurafenib induced mislocalization of desmosomal proteins (**Figure 2B**). Compared to DMSO, NHEKs treated with B-RAF inhibitors displayed a larger amount of DSG3 and PG in the cytoplasm rather than concentrated at intercellular borders, where they are needed to anchor keratin filaments and mediate strong adhesion between neighboring keratinocytes. This was reflected in line-scans across images of plakoglobin; peaks reflecting high concentration of the desmosomal protein at intercellular borders were markedly diminished in vemurafenib- or dabrafenib-treated cells (**Figure 2C**).

**Figure 2.**
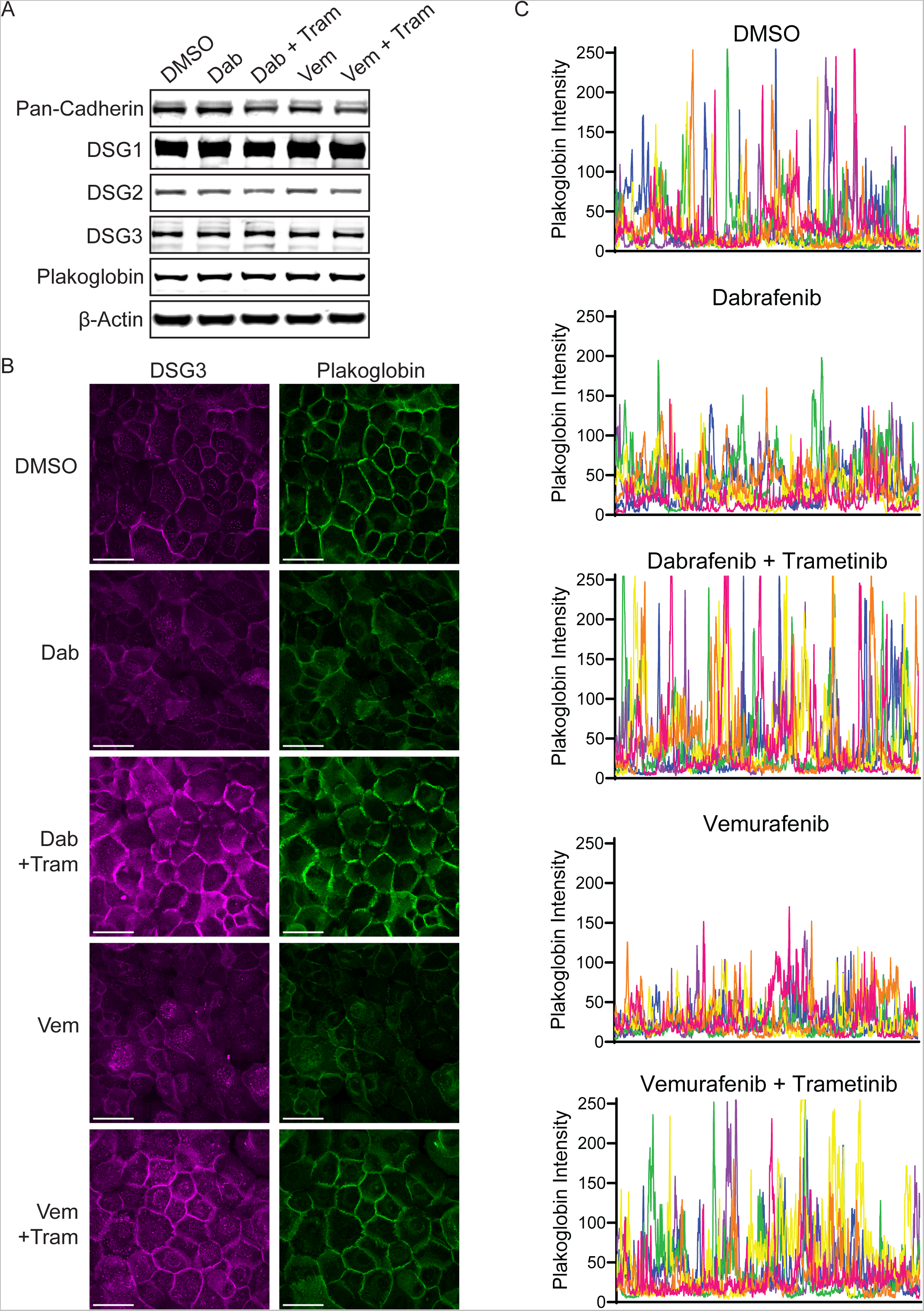
B-RAF inhibition disrupts desmosomal protein localization in epidermal keratinocytes. **(A)** Immunoblot of classical cadherins (Pan-Cad), desmosomal cadherins (DSG1, DSG2, DSG3), and plakoglobin in lysates from NHEKs treated with dabrafenib (Dab) or Vemurafenib (Vem) +/-trametinib (Tram) for 24 h; β-actin is a loading control. **(B)** Confocal immunofluorescence images of DSG3 (magenta) and plakoglobin (green) in NHEKs treated with the indicated compounds for 24 h; scale bar = 50 µm. **(C)** Line-scans were performed in a blinded manner across the entire field of N=6 confocal microscopy images (individually colored pink, orange, yellow, green, blue, or purple) for each drug condition; graphs depict plakoglobin fluorescence intensity of each pixel across the entire field of view with the largest peaks occurring as the line-scan crosses properly formed cell-cell junctions.

Using a mechanical dissociation assay validated for measuring intercellular adhesive strength via desmosomes in keratinocyte sheets (38), we demonstrated that the disruption of desmosomal organization noted in B-RAF-inhibited NHEKs translated into marked weakening of intercellular adhesion. NHEK monolayers treated with either dabrafenib or vemurafenib exhibited a significant increase in the number of fragments generated upon mechanical stress, reflecting reduced intercellular adhesive strength (**Figure 3A-B**). Together, these results indicate that B-RAF inhibitors can induce GD pathology through impaired localization of adhesive proteins, which weakens desmosomes to cause the severing of cell-cell junctions (acantholysis) in epidermal keratinocytes seen in GD biopsies, which manifests clinically as skin erosions and crusting.

**Figure 3.**
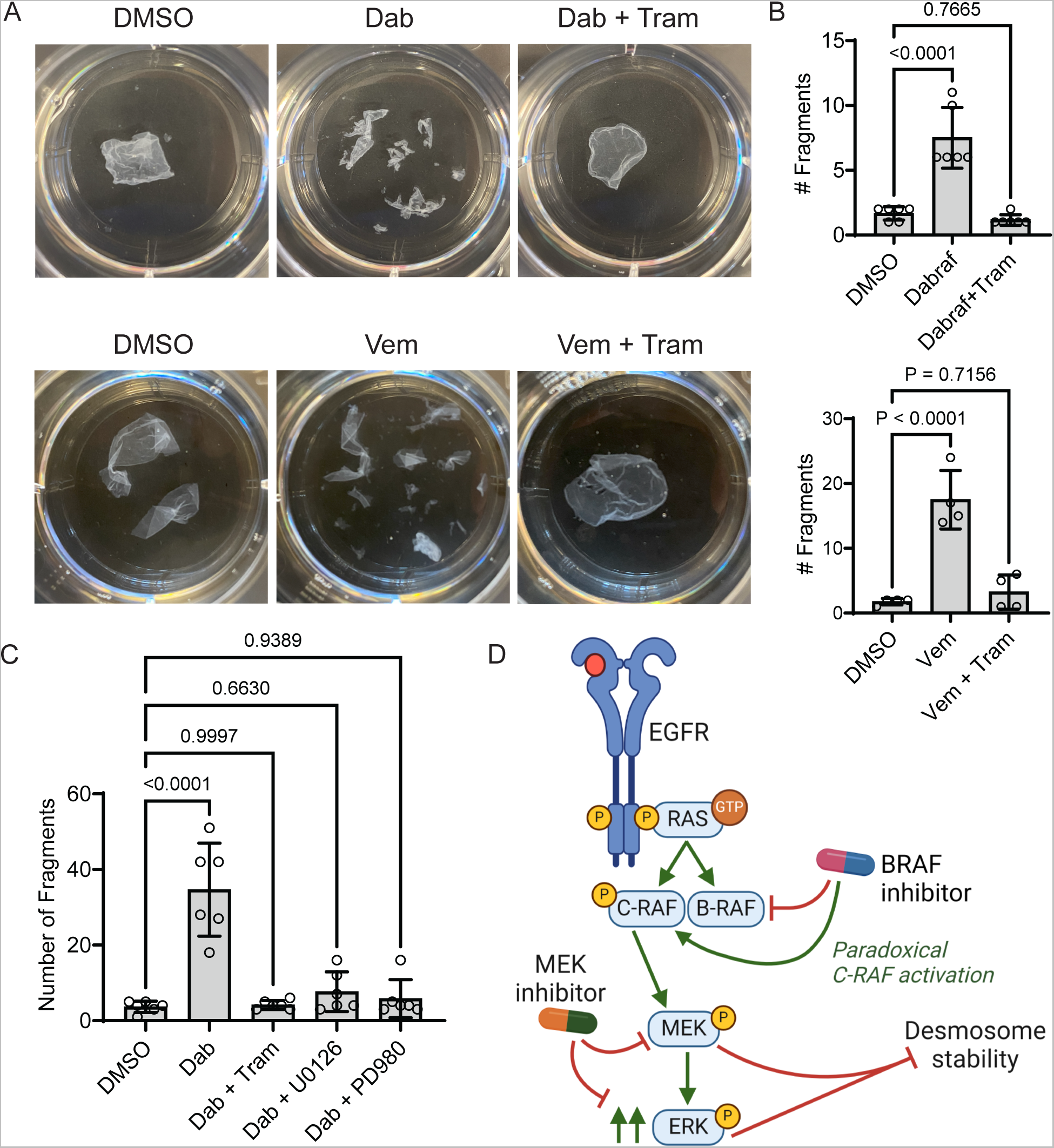
MEK suppression reverses B-RAF inhibitor-induced weakening of intercellular adhesion. **(A)** Mechanical dissociation assay of confluent monolayers from NHEKs cultured with the indicated compounds for 24 h; representative images of fragmented monolayers transferred into 6-well cell culture plates are shown. (**B**) Bar graphs display the mean ± SD of the number of fragments from drug-treated monolayers with individual data points plotted for N=6 (Dab) or N=4 (Vem) biological replicates; P-values from one-way ANOVA with Dunnett adjustment for multiple comparisons to control cells. **(C)** Bar graph displays the mean ± SD of the number of fragments from drug-treated monolayers with individual data points plotted for N=6 biological replicates; P-values are from one-way ANOVA with Dunnett adjustment for multiple comparisons to control cells. **(D)** Diagram depicts desmosome destabilization by MAP kinase pathway dysregulation; B-RAF inhibitors paradoxically activate C-RAF along with MEK and ERK downstream, which inhibits desmosome stability to cause Grover disease pathology, an effect overcome by MEK inhibitors.

### MEK suppression reverses B-RAF inhibitor-induced weakening of intercellular adhesion

Given that co-treatment of patients with MEK inhibitors prevented B-RAF inhibitor-induced GD in retrospective clinical studies (23), we hypothesized that MEK inhibitors would rescue cell-cell junctions in dabrafenib- or vemurafenib-treated NHEKs as a model of drug-induced GD. We found trametinib (Tram), a selective MEK inhibitor FDA-approved for *BRAF* mutant cancers (17, 39), greatly enhanced the localization of desmosomal proteins to cell-cell junctions in NHEKs despite treatment with dabrafenib or vemurafenib (**Figure 2B-C**).

Importantly, this rescue of cell-cell junctions in NHEKs translated into increased intercellular adhesive strength. While NHEK monolayers readily fragmented upon exposure to dabrafenib or vemurafenib, co-treatment with trametinib overcame the effect of B-RAF inhibitors, restoring the integrity of keratinocyte sheets to the level of control cultures treated with drug vehicle (**Figure 3A-B**). Confirming the specificity of our findings, we tested three additional MEK inhibitors (U0126, PD98059, cobimetinib), which comparably rescued cell cohesion in NHEK monolayers treated with a B-RAF inhibitor (**Figure 3C** and **Supplementary Figure S1**). In contrast, blocking other MAP kinases did not rescue intercellular adhesion in B-RAF-inhibited keratinocytes; p38 inhibition failed to restore the integrity of monolayers, while JNK inhibition actually led to an *increase* in fragmentation (**Supplementary Figure S1**). These data support the specificity of the therapeutic effect of dampening ERK activity to reverse B-RAF inhibitor-induced activation of MEK and resultant disruption of desmosomal adhesion in keratinocytes (**Figure 3D**).

### B-RAF inhibitors reversibly disrupt cell-cell junctions in organotypic human epidermis

To test the effects of B-RAF inhibitors in a 3-D human tissue context, we grew NHEKs as organotypic epidermis, which replicates fully differentiated epidermal morphology within a week (34), then treated mature cultures with vemurafenib for 48 hrs. Similar to our ImF results showing impaired desmosome organization in B-RAF-inhibited keratinocyte monolayers (**Figure 2B**), immunostaining of desmosomal components in organotypic epidermis treated with vemurafenib revealed impaired concentration of DSG1 and PG at cell-cell junctions (**Figure 4A-B**), These data are also consistent with prior work that showed mislocalization of desmosomal proteins, including DSG1 and PG, within skin biopsies of GD (24).

**Figure 4.**
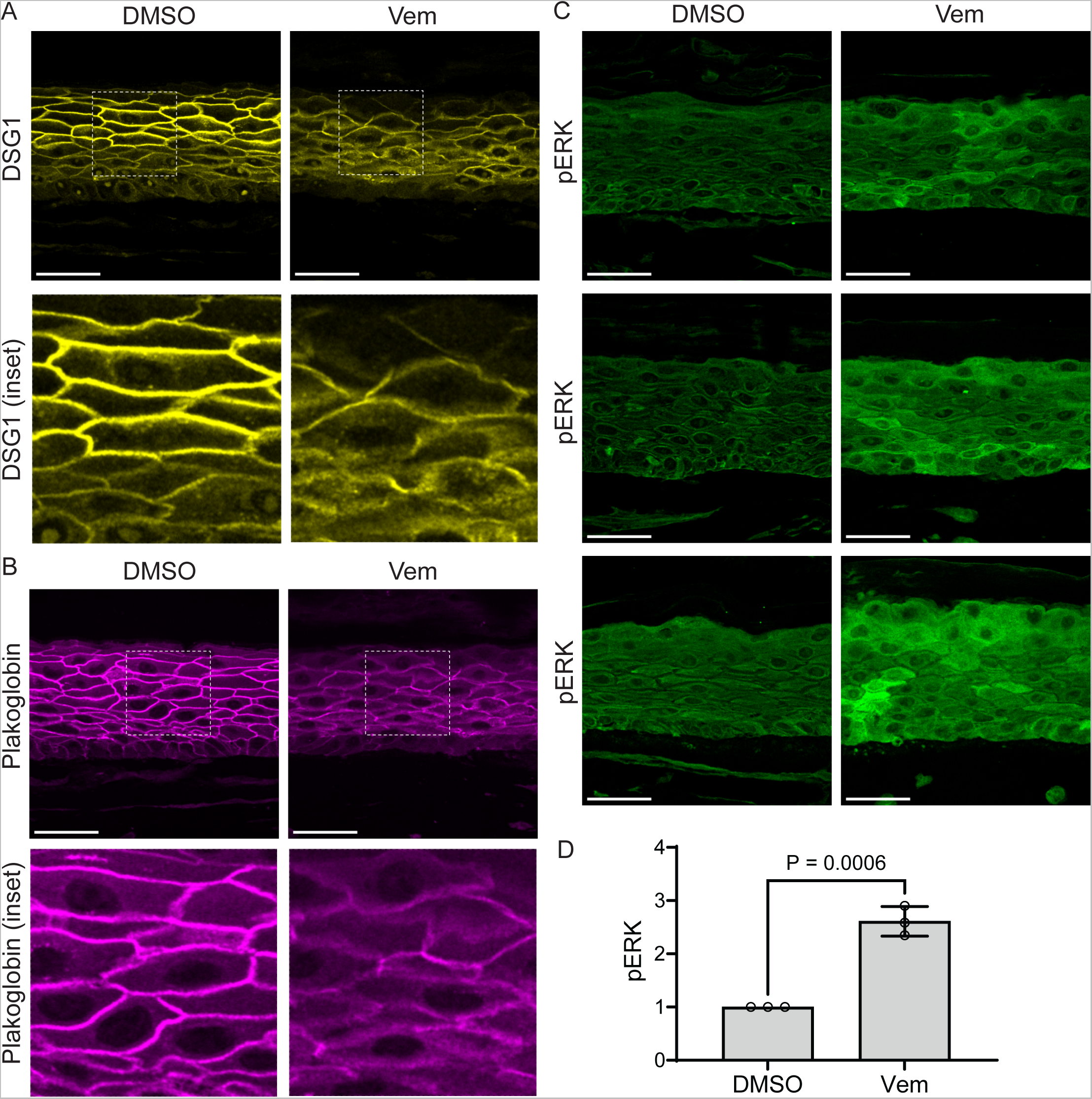
B-RAF inhibition is sufficient to disrupt cell-cell junctions and hyperactivate ERK in organotypic human epidermis. **(A)** Immunostaining of DSG1 (yellow) and **(B)** plakoglobin (magenta) in tissue cross-sections from organotypic epidermal cultures after 48 h of treatment with DMSO or vemurafenib (Vem), which disrupted desmosomal protein localization to cell-cell borders; scale bar = 50 µm; insets magnified below. **(C)** Immunostaining of pERK (green) in tissue cross-sections from epidermal cultures treated with DMSO vs. vemurafenib; scale bar = 50 µm. **(D)** Quantification of epidermal immunostaining of pERK in cross-sections of DMSO-vs. vemurafenib-treated cultures; bar graph displays the mean (individual values plotted) ± SD of pERK intensity from ≥60 images from N=3 biological replicates for each drug.

Despite this marked alteration in cell-cell junction morphology, we did not find areas of complete acantholysis in vemurafenib-treated tissue sections, which is likely due to reduced shear mechanical forces on desmosomes within the *in vitro* tissue model compared to epidermis *in vivo*. Consistent with this, our mechanical dissociation assay required the application of shear stress to sever weakened intercellular adhesions and induce fragmentation of vemurafenib-treated keratinocyte monolayers (**Figure 3A-B**). Similar to our results showing increased ERK activation by vemurafenib in keratinocyte monolayers (**Figure 1**), immunostaining of tissue sections from vemurafenib-treated epidermal cultures revealed a significant increase in pERK levels (**Figure 4C-D**). This led us to investigate whether vemurafenib treatment induced ERK activation in patients who developed Grover disease as a side effect of therapy.

### Biopsies of B-RAF inhibitor-induced Grover disease show ERK hyperactivation

Since GD is not a genetic disorder that can be easily replicated using knockout cells or mice, we established a model of drug-induced GD using B-RAF inhibitors that have been robustly linked in clinical studies to inducing specific GD pathology in patients (14, 15). To validate the findings from our *in vitro* model of drug-induced GD, we aimed to assess ERK activation levels in skin biopsies from patients treated with B-RAF inhibitor monotherapy. We obtained fixed tissue sections of skin biopsies from a de-identified cohort of five patients treated with vemurafenib, who developed a cutaneous eruption with pathologic features diagnosed as drug-induced GD by a board-certified dermatopathologist (**Figure 5A**).

**Figure 5.**
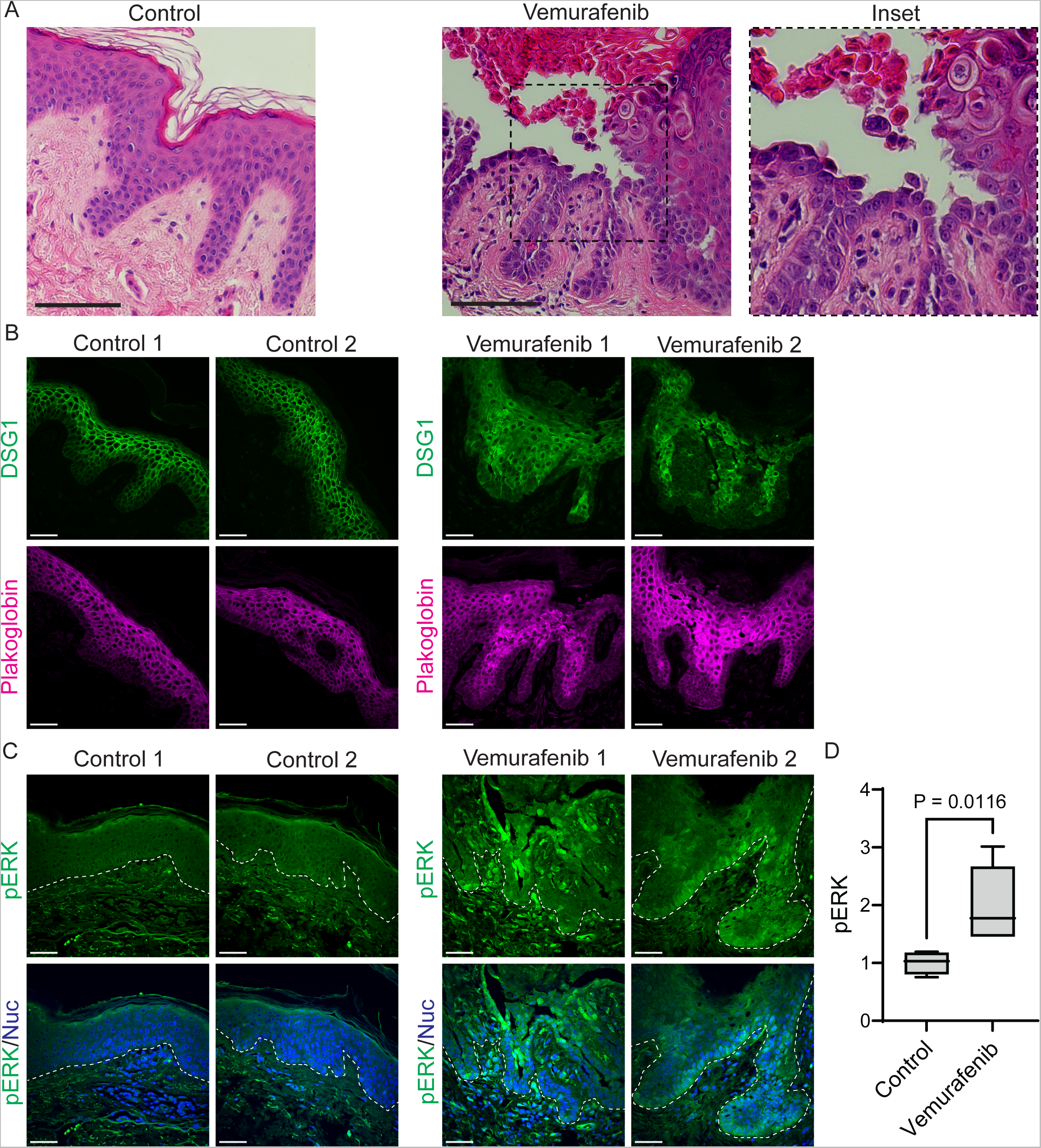
Biopsies of B-RAF inhibitor-induced Grover disease show epidermal hyperactivation of ERK and disruption of desmosomal proteins. **(A)** H&E-stained cross-sections of punch biopsies from the skin of a control donor versus a patient diagnosed with vemurafenib-induced Grover disease, which demonstrates aberrant cornification (dyskeratosis) and epidermal splitting between keratinocytes (magnified in inset); scale bar = 100 µm. **(B)** Immunostaining of DSG1 and plakoglobin in tissue cross-sections from patient biopsies; images shown are from 2 control and 2 patients with drug-induced Grover disease and are representative of N=5 patients in each group; scale bar = 50 µm. **(C)** Immunostaining of pERK and Hoechst to stain nuclei (Nuc) in tissue cross-sections from patient biopsies; images shown are from 2 control and 2 patients with drug-induced Grover disease and are representative of N=5 patients in each group; dashed line marks bottom of the epidermis; scale bar = 50 µm. **(D)** Quantification of epidermal immunostaining of pERK in cross-sections of control biopsies versus in lesions of drug-induced Grover disease; pERK intensity data for each group are depicted as a box plot of the 25^th^-75^th^ percentile with a line at the median from N=5 control vs. N=5 Grover disease biopsies; control mean normalized to 1; P-value from Student’s t-test.

Immunostaining biopsies of vemurafenib-induced GD skin lesions compared to normal control skin, we found disruption in the localization of both DSG1 and PG at cell-cell borders in the epidermis (**Figure 5B**). These data are consistent with prior studies of GD (24), which demonstrated a breakdown of desmosomal adhesion within areas of keratinocyte acantholysis typical of GD lesions. Moreover, we found a significant increase in pERK levels within the epidermal lesions of vemurafenib-related GD (**Figure 5C-D**). These data confirm in patients treated with vemurafenib monotherapy that B-RAF inhibition is associated with a paradoxical increase in ERK activation within skin lesions that exhibit loss of cell-cell adhesion and were diagnosed as drug-induced GD, thus directly implicating ERK signaling in GD pathogenesis. Our results explain why the addition of MEK inhibitors like trametinib, which suppress ERK over-activation, can eliminate GD as a side effect of B-RAF inhibitors.

### Idiopathic Grover disease lesions exhibit increased ERK activation

Based on our results showing that B-RAF inhibition causes over-activation of ERK in human keratinocytes, organotypic epidermis, and in patients with drug-induced GD, we proposed that this same mechanism could drive the pathology of spontaneous GD, a more common dermatologic condition that is considered idiopathic and currently lacks any FDA-approved therapy. To test this, we obtained tissue sections from 17 skin biopsies from a de-identified cohort of patients with typical pathologic features of spontaneous GD (**Figure 6A**) and performed immunostaining for both intercellular junctions and ERK activation.

**Figure 6.**
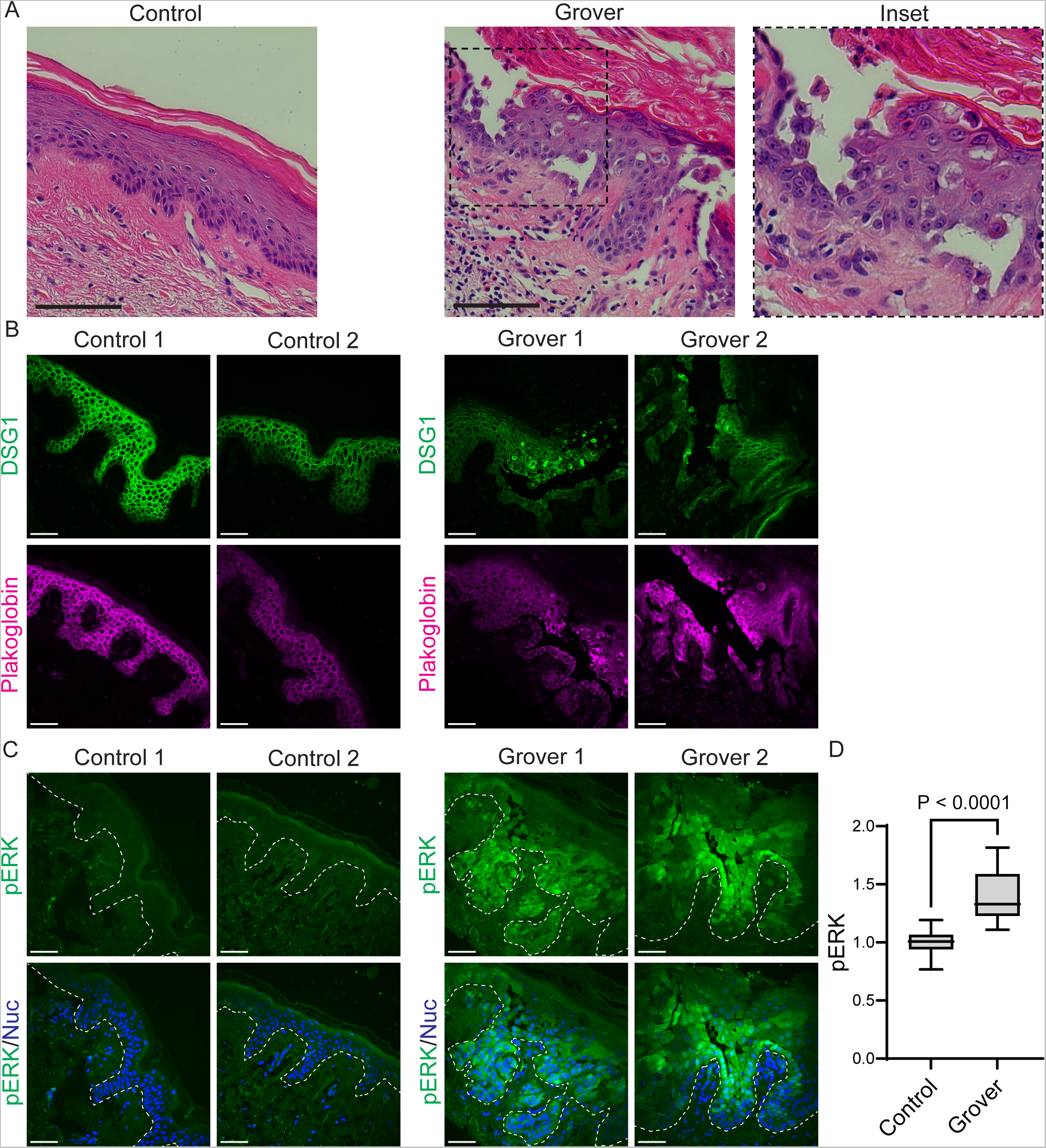
Idiopathic Grover disease biopsies exhibit ERK hyperactivation along with desmosomal disruption. **(A)** H&E-stained cross-sections of punch biopsies from the skin of a control donor versus a patient with idiopathic Grover disease, which demonstrates aberrant cornification (dyskeratosis) with retention of nuclei in the cornified layers and loss of keratinocyte cohesion (magnified in inset); scale bar = 100 µm. **(B)** Immunostaining of DSG1 and plakoglobin in tissue cross-sections from patient biopsies; images shown are from 2 control and 2 Grover disease patients and are representative of N=17 patients in each group; scale bar = 50 µm. **(C)** Immunostaining of pERK and Hoechst to stain nuclei (Nuc) in tissue cross-sections from patient biopsies; images shown are from 2 control and 2 Grover disease patients and are representative of N=17 patients in each group; dashed line marks bottom of the epidermis; scale bar = 50 µm. **(D)** Quantification of epidermal immunostaining of pERK in cross-sections of biopsies from control versus Grover disease lesions; pERK intensity data for each group are depicted as a box plot of the 25^th^-75^th^ percentile with a line at the median from N=17 control vs. N=17 Grover disease biopsies; control mean normalized to 1; P-value from Student’s t-test.

Compared to biopsies of control skin, staining of GD biopsies revealed clear disruption of cell-cell junctions between epidermal keratinocytes. Within lesional areas, DSG1 and PG were nearly completely internalized within the cytoplasm and collapsed around the nucleus of epidermal keratinocytes rather than being localized to intercellular borders (**Figure 6B**) Importantly, we also found a concomitant increase in ERK phosphorylation within GD lesions (**Figure 6C-D**). Together, our results implicate overactivation of the MAP kinase pathway via ERK in the pathogenesis of both drug-induced and idiopathic GD and underscore the potential therapeutic value of MEK inhibitors (several of which are already cleared for clinical use) for this dermatologic disorder that is in need of new treatment strategies (**Graphical Abstract**).

## DISCUSSION

Molecular therapies have completely changed the treatment landscape and prognosis for malignancies driven by specific mutations, such as *BRAF* V600E in melanoma (40). Despite being highly selective for their targets, inhibitors of the MAP kinase signaling pathway have had unanticipated side effects, including frequent cutaneous adverse events that reduced drug safety and tolerability, impaired quality of life, and even disqualified patients from trials (41, 42). As a silver lining, off-target effects of a drug can provide insight into the pathogenesis of other diseases. In clinical trials of B-RAF inhibitor monotherapy for cancer, investigators reported an unexpectedly common skin eruption with biopsy features diagnostic of GD (14, 15); subsequent analysis indicated GD was seen in 42.9% and 38.9% of patients treated with dabrafenib or vemurafenib, respectively (23). However, it was unclear how B-RAF blockade replicated the specific pathologic findings of a rare skin blistering disorder that had not previously been linked to this signaling pathway. We used multiple *in vitro* assays to demonstrate that B-RAF inhibitors are sufficient to increase ERK activation, which disrupted desmosomal adhesion in human keratinocytes and organotypic epidermis, thus explaining the loss of tissue integrity typical of GD pathology.

Intriguingly, the pathologic features of GD can be identical to Darier disease (9, 10, 24), a genetic disorder linked to mutation of a calcium ATPase (SERCA2) embedded in the endoplasmic reticulum (11). Up to now, it remained unclear why these two disorders would exhibit such similar findings in biopsies that pathologists cannot consistently distinguish the diagnoses without additional clinical information, such as whether the eruption is known to be hereditary (Darier disease) or spontaneous (Grover disease). Our recent work established an *in vitro* model of Darier disease and demonstrated that deficiency or chemical inhibition of SERCA2 induced hyperactivation of ERK (22), which we report here as a driver of GD pathology. Further linking these disorders with distinct origins (inherited vs. acquired), some cases of GD were recently found to harbor acquired mutations in the Darier disease-linked gene *ATP2A2* encoding SERCA2, a major regulator of calcium, which can activate ERK signaling in keratinocytes (43). The convergence of these two disorders upon the same signaling pathway explains the similarity of their pathologic features. Moreover, recent RNA sequencing of skin biopsies from patients with Grover or Darier disease identified an overlapping transcriptional signature that pointed to dysregulation of serum response factor and the actin cytoskeleton (44), both of which are modulated by MEK and ERK (45-47).

While molecular therapies have revolutionized the treatment of common inflammatory dermatologic conditions using monoclonal antibodies against cytokines or selective kinase inhibitors (48-51), targeted therapeutics remain elusive for rare blistering disorders. Current therapy of GD is based on case series and expert opinion as it lacks any treatment proven in prospective trials (7, 8, 52). Retinoids, which regulate the differentiation of keratinocytes, have been used in observational studies for GD (7, 8, 53-55), but they confer a high risk of toxicity with long-term use and are potent teratogens (56, 57). Other conventional, but off-label, therapies for GD include topical corticosteroids and repeated narrow-band ultraviolet light treatment, which is burdensome for patients (7, 8). More recently, dupilumab has been reported as an off-label treatment for refractory GD (58-60), though its blockade of interleukin-4 and -13 receptors may target secondary pruritus and inflammation rather than the primary pathogenic drivers of GD. Our findings suggest MEK inhibitors, which are FDA-approved for oral administration for multiple *BRAF*-driven cancers (17, 61-65), could be therapeutic for GD as well as Darier disease. Moreover, MEK inhibitors can be delivered topically as shown by successful use of compounded trametinib to treat a cutaneous histiocyte proliferation driven by MAP kinase overactivation (66). Topical MEK inhibition could obviate side effects from systemic use (23, 67, 68) and their delivery is likely to be efficient for blistering disorders like GD given that their pathology lies in the epidermis and already compromises the cutaneous barrier.

In conclusion, our results from cellular and organotypic models identified ERK as a key driver of GD pathology and demonstrate that MEK inhibition was sufficient to restore desmosomal organization and rescue intercellular adhesion in keratinocytes with hyperactive ERK signaling. Moreover, our finding of ERK hyperactivation in skin lesions from patients with both B-RAF inhibitor-induced and idiopathic GD substantiate MAPK signaling modulation as a viable strategy for GD treatment. The existence of multiple FDA-approved agents targeting MEK, in particular, further supports their feasibility for clinical trials for GD. While the therapeutic potential of MEK inhibitors for GD in patients has not yet been tested, data from multiple clinical trials revealed that adding trametinib to B-RAF inhibitors eliminated drug-induced GD (23), which provides excellent rationale for clinical studies to evaluate MEK inhibitors for treatment of idiopathic GD.

## METHODS

### Sex as a biological variable

Our *in vitro* studies examined available human primary and immortalized keratinocytes from males, but our results were validated using skin biopsy specimens from both male and female patients, which showed similar findings.

### Reagents

Inhibitors of MEK including trametinib (Cat. #62206), U0126 (Cat. #9903), PD98059 (Cat. #9900), PD184352 (Cat. #12147) plus inhibitors of B-RAF including dabrafenib (Cat. #91942) and vemurafenib (Cat. #17531) were from Cell Signaling Technology. Rabbit antibodies recognizing phospho-ERK1/2 (D13.14.4E; Cat. #4370), pan-cadherin (Cat. #4068), plakoglobin (Cat. #75550) and mouse anti-ERK1/2 (L34F12; Cat. #4696) came from Cell Signaling Technology. Rabbit anti-KRT10 (Cat. #ab76318) came from Abcam. Mouse anti-DSG1 (Cat. #sc-137164), anti-DSG2 (Cat. #sc-80663), anti-DSG3 (Cat. #sc-53487), anti-plakoglobin (Cat. #sc-514115), anti-GAPDH (Cat. #sc-47724), and anti-β-Actin (C4; Cat. #sc-47778) came from Santa Cruz Biotechnologies. Chicken anti-plakoglobin (#1408) was a gift from Dr. Kathleen Green (Northwestern Univ., Chicago, IL, USA). Secondary antibodies for fluorescent immunoblotting, including IRDye 800CW goat anti-rabbit IgG (Cat. #926-32211) and IRDye 680RD goat anti-mouse IgG (Cat. #926-68070), came from LI-COR Biosciences. For cell and tissue staining, fluorescent secondary antibodies came from Thermo-Fisher: Goat anti-mouse IgG AlexaFluor-405 (Cat. #A31553), AlexaFluor-488 (Cat. #A11001), AlexaFluor-594 (Cat. #A11005), or AlexaFluor-633 (Cat. #A21050); goat anti-rabbit IgG AlexaFluor-405 (Cat. #A31556), AlexaFluor-488 (Cat. #A11008), AlexaFluor-594 (Cat. #A11012), or AlexaFluor-633 (Cat. #A21070). Hoechst 33342 came from Thermo-Fisher (Cat. #H1399). The ERK fluorescent biosensor (pLenti-CMV-Puro-DEST-ERK-KTR-mClover; Cat. #59150) was from Addgene.

### Cell culture

Normal human epidermal keratinocytes (NHEKs) were cultured in Medium 154 adjusted to 0.07 mM CaCl^2^ (Thermo-Fisher Cat. #M154CF500) plus 1x human keratinocyte growth supplement (Thermo-Fisher Cat. #S0015) and 1x gentamicin/amphotericin (Thermo-Fisher Cat. #R01510).

J2-3T3 immortalized murine fibroblasts were cultured in complete Dulbecco’s Modified Eagle Medium (DMEM) (Thermo-Fisher Cat. #11965092) with a final concentration of 10% FBS (Hyclone, Fisher Scientific Cat. #SH3039603), 2 mM GlutaMAX (Thermo-Fisher Cat. #35050061), 100 U/mL penicillin, and 100 μg/mL streptomycin.

All cell lines were grown at 37°C in 5% CO^2^ within an air-jacketed, humidified incubator. Cells were cultured on sterile tissue culture-treated dishes and passaged using 0.25% Trypsin-EDTA (Thermo-Fisher Cat. #15400054) while sub-confluent.

### Organotypic epidermal culture

Organotypic human epidermal “raft cultures” were grown as described (34, 69). Cultures were differentiated in E-medium, containing a 3:1 mixture of DMEM:Ham’s F12 (Thermo-Fisher Cat. #11765054) along with 10% FBS, 180 µM adenine (Sigma Cat. #A2786), 0.4 µg/mL hydrocortisone (Sigma, Cat. #H0888), 5 µg/mL human insulin (Sigma Cat. #91077C), 0.1 nM cholera toxin (Sigma, Cat. #C8052), 5 µg/mL apo-transferrin (Sigma Cat. #T1147), plus 1.36 ng/mL 3,3′,5-tri-iodo-L-thyronine (Sigma Cat. #T6397).

J2-3T3 murine fibroblasts were suspended in collagen matrix rafts in transwells (Corning Cat. #353091). Per raft, 1 x 10^6^ fibroblasts were resuspended in 10% of the final desired volume of sterile reconstitution buffer (1.1 g of NaHCO^3^ and 2.39 g of HEPES in 50 mL 0.05 N NaOH), to which was added 10% of the final desired volume of 10X DMEM (Sigma-Aldrich Cat. #D2429). After thoroughly mixing the cells by pipetting, high-concentration rat tail collagen I (Corning Cat. #CB354249) was added (to a final concentration of 4 mg/mL) and the slurry was supplemented with sterile diH2O to dilute the solution to the final desired volume, 2 mL total per raft. As needed, 0.05 N NaOH was added dropwise until the pH reached ∼7 based on the indicator phenol red.

The slurry of collagen and fibroblasts was inverted to mix, then was 2 mL was pipetted into each transwell insert suspended within a deep 6-well cell culture plate (Corning Cat. #08-774-183). The rafts were allowed to polymerize for one hour at 37°C, then were submerged in 16 mL complete DMEM and allowed to incubate overnight at 37°C.

The next day, keratinocyte cultures were trypsinized then resuspended in E-medium containing EGF (5 ng/mL) to final concentration of 0.75 x 10^6^ cells/mL (in a final volume of 2 mL per organotypic culture). The DMEM was removed from the upper and lower transwell chambers by aspiration, then 2 mL (1.5 x 10^6^ cells) of resuspended keratinocytes were pipetted on top of each raft. E-medium plus 5 ng/mL EGF was added to both the top and bottom transwell chambers to submerge the raft, then the cultures were incubated overnight at 37°C. After waiting 24 hours, the E-medium was removed from both the top and bottom chambers. To initiate stratification, keratinocytes were placed at an air-liquid interface by adding E-medium (lacking EGF) to only the bottom chamber until reaching the bottom of the raft. Organotypic cultures were grown for 8-12 days at 37°C with replacement of E-medium in the bottom chamber every 2 days. Chemical inhibitors or vehicle control (DMSO) were diluted in the bottom chamber E-medium. The concentration of inhibitors used were: dabrafenib (1 µM), vemurafenib (10 µM), trametinib (1 µM).

To perform histology, the entire transwell and raft culture were submerged in 10% neutral-buffered formalin (Fisher Scientific Cat. #22-026-435) within a 6-well cell culture plate for at least 24h. Organotypic epidermis was processed for histologic examination plus H&E staining by the Fred Hutchinson Cancer Center Experimental Histopathology Core.

### Immunoblotting

NHEKs seeded at 1 x 10^6^ cells per well of 6-well cell culture dishes were grown in M154 until reaching confluence, then were switched into E-medium with vehicle control (DMSO) or inhibitors as follows: Dabrafenib (1 µM), vemurafenib (10 µM), trametinib (1 µM), cobimetinib (1 µM), selumetinib (1 µM), U0126 (10 µM), PD98059 (20 µM), SB203580 (10 µM), or SP600125 (25 µM). After 24-48 h, whole-cell lysates were made after washing cells in PBS then applying urea sample buffer [8 M Urea, 1% SDS, 10% glycerol, 0.0005% pyronin-Y, 5% β-mercaptoethanol, 60 mM Tris, pH 6.8] for 10 min. Lysates homogenization was performed using a microtip probe sonicator (Fisher Scientific).

Protein lysates were loaded into NuPAGE 12% Bis-Tris Gels (Thermo-Fisher Cat. #NP0343BOX) then separated by electrophoresis in NuPAGE MES SDS Running Buffer (Thermo-Fisher Cat. #NP0002). Proteins were then transferred onto Immobilon-FL membrane (Millipore Cat. #IPFL85R) using transfer buffer (25 mM Tris, 192 mM glycine, 20% (v/v) methanol) for 60 minutes at 50 V. Membranes were blocked for 60 min at room temperature in Intercept tris-buffered saline (TBS) blocking buffer (LI-COR). Membranes were probed at 4°C overnight in primary antibodies in Intercept TBS blocking buffer (LI-COR). Blots were washed in 1x TBS containing 0.1% (v/v) Tween-20 (TBS-T) three times, then were incubated 1 h at room temperature in Intercept TBS blocking buffer containing IRDye 800CW goat anti-rabbit IgG and/or IRDye 680RD goat anti-mouse IgG (LI-COR) at 1:10,000. Blots were washed in TBS-T three times, then protein bands were visualized on an Odyssey M Imaging System (LI-COR).

### Fluorescent cell staining

Keratinocytes were seeded in 35 mm glass-bottom cell culture dishes (MatTek Cat. #P35G-1.5-20-C) and grown to confluency. For staining desmosomal proteins and keratins, cells were fixed at -20°C for 2 min using ice-cold 100% methanol, dried, then re-hydrated in PBS. For all other staining, cells were fixed using 4% paraformaldehyde at 37°C for 10 min. Fixed cells were then incubated for 30 min at 37°C in blocking buffer [0.5% (w/v) bovine serum albumin (BSA, Sigma Cat. #A9647), 10% (w/v) normal goat serum (NGS, Sigma Cat. # G9023) in PBS]. Cells were washed once in PBS, then primary antibodies diluted in 0.5% (w/v) BSA in PBS were applied to the cells overnight at 4°C. Dilutions of primary antibodies were: mouse anti-DSG3 (1:50) and rabbit anti-plakoglobin (1:50). Cells were rinsed in PBS 3 times, then were incubated for 60 min at 37°C in secondary antibodies diluted 1:300 (+/-Hoechst at 1:500) in 0.5% (w/v) BSA in PBS. Cells were rinsed three times in PBS and held in PBS for confocal microscopy.

### Tissue processing and histologic analysis

Formalin-fixed paraffin-embedded tissue cross-sections from human organotypic epidermis or skin biopsies underwent standard histology processing followed by hematoxylin and eosin (H&E) staining. H&E-stained tissues were imaged using a 40X long working distance, achromatic, phase-contrast objective on the EVOS FL microscope (Thermo-Fisher). H&E images were obtained on the EVOS high-sensitivity embedded interline CCD color camera.

### Fluorescent tissue staining

Formalin-fixed paraffin-embedded tissue cross-sections on glass slides were baked for 2 h at 65°C. Baked slides were immersed for 3 min in each of: 3 baths of xylenes (Fisher Cat. #X3P), 3 baths of 95% ethanol, 70% ethanol, and 3 baths of PBS. Slides were then heated to 95°C for 15 min in antigen retrieval buffer [0.1 M sodium citrate (pH 6.0) with 0.05% (v/v) Tween-20]. After cooling to room temperature, slides were rinsed in PBS. Tissue sections were encircled using a PAP pen hydrophobic barrier. Tissue sections were incubated in a humidified chamber for 30 min at 37°C in blocking buffer [0.5% (w/v) BSA, 10% (v/v) NGS in PBS]. Slides were rinsed in 3 PBS baths for 3 min each, then were incubated overnight at 4°C in a humidified chamber in primary antibodies diluted in 0.5% (w/v) BSA in PBS. Primary antibodies were diluted as follows: mouse anti-desmoglein 1 (1:50), mouse anti-ERK (1:400), rabbit anti-phospho-ERK (1:400), rabbit anti-cytokeratin 10 (1:3000), rabbit anti-plakoglobin (1:100), or chicken anti-plakoglobin (1:1000). Slides were then washed for 3 min in each of 3 baths of PBS then were incubated for 60 min at 37°C in a humidified chamber in secondary antibodies diluted at 1:300 (+/-Hoechst at 1:500) in 0.5% (w/v) BSA in PBS. Slides were washed for 3 min each of 3 baths of PBS. Finally, a drop of Prolong Gold (Thermo-Fisher Cat. #P36934) was applied over the tissue sections along with a #1.5 glass coverslip. After drying, slides were imaged using confocal microscopy.

### Confocal fluorescence microscopy

A Hamamatsu ORCA-FusionBT sCMOS camera and Yokogawa W1 spinning-disk confocal (SDC) system on a Nikon Ti2 microscope were used to acquire images. Samples were illuminated with laser excitation lines (405, 488, 561, 640 nm) and emitted fluorescence signal was captured through a 60x 1.2 NA water objective (Nikon) and standard filters.

### ERK biosensor imaging

For live imaging of the ERK biosensor, NHEKs were transduced with ERK-KTR-mClover (Addgene Cat. #59150). HEK293FT cells were grown in complete DMEM, then transfected with 4 μg pLenti-CMV-PuroDEST-ERK-KTR-mClover DNA plus 12 μL FuGENE 6 (Promega Cat. #E2691) in 800 μL of Opti-MEM (Thermo-Fisher Cat. #31985070), which was added to the cells and left overnight. Lentiviral supernatants were collected the next day and polybrene (Sigma Cat. #H9268) was added (4 μg/mL). M154 was removed from NHEKs and replaced with viral medium for 1 h at 37°C. After washing in PBS, M154 was replaced and cells were expanded in culture.

ERK-KTR-mClover-transduced cells were seeded into 35 mm glass-bottom dishes in low-calcium medium (0.31 mM) and grown to confluency. Cells were then exposed to high-calcium (1.3 mM) for 24 h, then imaged by SDC microscopy. Using Fiji, the nuclear region was encircled with the polygon tool and the “Measure” function was used to calculate the mean fluorescence intensity; after using the “Cut” function to remove the nuclear region, then remaining cytoplasmic region was encircled with the polygon tool and the “Measure” function was used to calculate the mean fluorescence intensity. The ERK activity index was calculated as the ratio of the cytoplasmic integrated fluorescence intensity (active) to the nuclear integrated fluorescence intensity.

### Fluorescent tissue staining quantification

Immunostained tissue sections images were obtained using SDC microscopy as above and were quantified using Fiji. Fluorescence intensity was measured on non-visibly labeled images of immunostained tissue cross-sections. The epidermis was circumscribed using the Polygon tool, then the “Measure” function was utilized to calculate the mean fluorescence intensity of the same encircled region for each laser channel. Fluorescence intensity mean from each image was averaged across all non-overlapping high-powered fields (hpf); the mean intensity of pooled control samples was normalized to 1.

### Monolayer mechanical dissociation assay

A mechanical dissociation assay was performed as described (70). Keratinocytes plated at 1 x 10^6^ cells per well of 6-well cell culture dishes were grown to confluence, then were switched into E-medium. Vehicle control (DMSO) was compared to chemical inhibitors used at the following concentrations: Dabrafenib (1 µM), vemurafenib (10 µM), trametinib (1 µM), U0126 (10 µM), PD98059 (20 µM), cobimetinib (1 µM), SP600125 (25 µM), SB203580 (10 µM). After 24h, treated monolayers were washed in PBS then incubated in 500 μL dispase (5 U/ml) in Hank’s balanced salt solution (Stemcell Technologies, Cat. #07913) at 37°C for 30 min. Then, 4.5 mL PBS was added to each well and released monolayers plus all liquid were transferred into 15 mL conical tubes; all tubes were placed in a rack and inverted together 1-10 times to apply mechanical stress. Fragmented monolayers were transferred into 6-well tissue culture plates then imaged using a 12-megapixel digital camera. Fragments were counted manually in images.

### Statistics

Prism version 9 (GraphPad) was used for statistical analyses and graphing. Each figure legend notes the included statistical parameters such as sample size, center definition, measures of dispersion, and statistical tests. Normality of datasets was tested using the D’Agostino-Pearson test. The means of two normally distributed groups were compared with a two-tailed unpaired Student’s t test. Means from greater than two normally distributed groups were compared with a one-way ordinary ANOVA with P-values adjusted for multiple comparisons. P-values <0.05 were deemed statistically significant. Each graph includes exact P-values.

### Study approval

The Penn Skin Biology and Diseases Resource-based Center (SBDRC) isolated normal human epidermal keratinocytes (NHEKs) from de-identified neonatal foreskins under a protocol (#808224) approved by the University of Pennsylvania Institutional Review Board (IRB). Sections of tissue from de-identified skin biopsies were procured by the SBDRC from a tissue bank under a protocol (#808225) approved by the University of Pennsylvania IRB. Use of de-identified tissues collected for clinical purposes, which otherwise would be discarded, was exempted for written informed consent by the IRB.

## Data availability

All underlying values for graphed data are available in the **Supporting Data Values** file.

## AUTHOR CONTRIBUTIONS

A.T. and S.A.Z. were deemed to have contributed equally to the hypotheses, experimental design, and data essential to the included studies. Conceptualization: S.A.Z., A.T., C.L.S. Data Curation: A.T., S.A.Z., E.Y.C., P.W.H., J.E.G., C.L.S. Formal analysis: A.T., S.A.Z., C.J.J., C.L.S. Investigation: S.A.Z., A.T., C.L.S. Resources: C.L.S., J.E.G. Supervision: C.L.S. Visualization: A.T., C.L.S. Validation: A.T., S.A.Z., E.Y.C., P.W.H., C.L.S. Writing (original draft): C.L.S. Writing (review and editing): C.L.S., A.T., S.A.Z., C.J.J., E.Y.C., P.W.H., J.E.G.

## ACKNOWLEDGEMENTS

The authors thank the Skin Biology and Diseases Resource-Based Centers at University of Pennsylvania (NIH P30-AR069589) and University of Michigan (NIH P30-AR075043) for histology services and procurement of archived biopsies and NHEKs. C.L.S. was supported by NIH K08-AR075846. A.T. was supported by NIH R03-AR082896, an Innovation Pilot Award from the University of Washington Institute for Stem Cell and Regenerative Medicine, and a grant from the Foundation for Ichthyosis and Related Skin Types. C.J.J. was supported by a Medical Scholars Research Fellowship from the Physician-Scientist Support Fund. P.W.H. was supported by NIH P30-AR075043. J.E.G. was supported by NIH P30-AR075043 and the Taubman Medical Research Institute. BioRender.com was used to make the Graphical abstract.

