## Supplemental Figure 1 for "ERK hyperactivation in epidermal keratinocytes impairs intercellular adhesion and drives Grover disease pathology"

Supplemental Figure S1

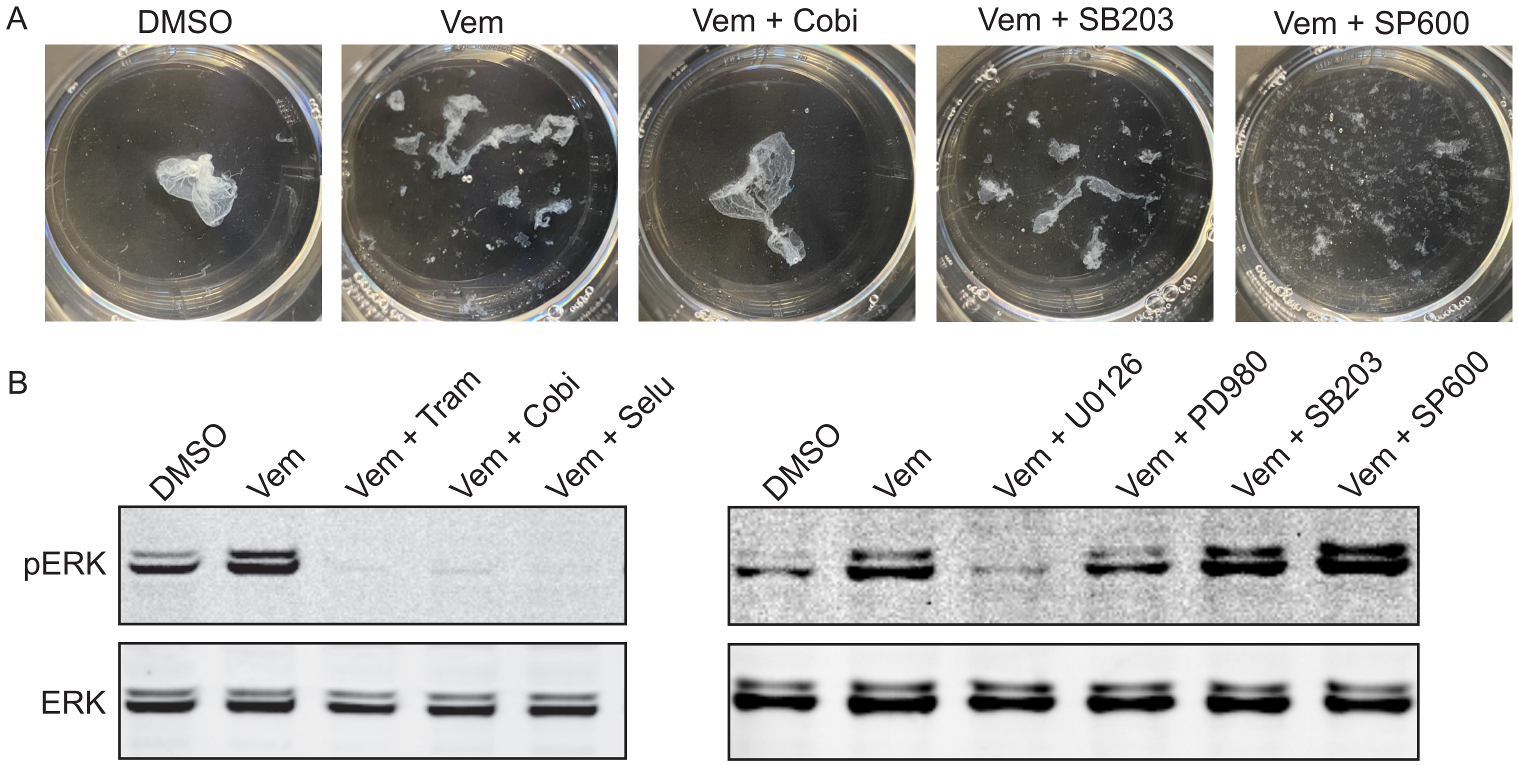


**Supplemental Figure S1 |** MEK inhibition dampens ERK hyper-activation and restores integrity of vemurafenib-treated keratinocyte sheets. **(A)** Images of drug-treated keratinocyte monolayers after a mechanical dissociation assay; representative images of fragmented cell sheets transferred into 6-well cell culture plates are shown. **(B)** Immunoblot of total and phosphorylated ERK (pERK) in lysates from NHEKs treated for 24 h with vemurafenib (Vem, 10 μM) +/- MEK inhibitors trametinib (Tram, 1 μM), cobimetinib (Cobi, 1 μM), selumetinib (Selu, 1 μM), U0126 (10 μM), PD98059 (PD980, 20 μM), a p38 inhibitor SB203580 (SB203, 10 μM), or the JNK inhibitor SP600125 (SP600, 25 μM).
